# Adipose gene expression profiles and their associations with adaptations in Northern Finncattle, Mirandesa cattle, Yakutian cattle and Holstein cattle

**DOI:** 10.1101/2023.12.21.572790

**Authors:** Daniil Ruvinskiy, Andreia Amaral, Melak Weldenegodguad, Mervi Honkatukia, Heli Lindeberg, Jaana Peippo, Päivi Soppela, Florian Stammler, Pekka Uimari, Catarina Ginja, Juha Kantanen, Kisun Pokharel

**Affiliations:** Natural Resources Institute Finland (Luke), Myllytie 1, FI-31600 Jokioinen, Finland; Escola de Ciência e Tecnologia, Universidade de Évora, Largo dos Colegiais, N° 2, 7004- 516 Évora, Portugal; Centro Interdisciplinar em Investigação em Sanidade Animal, Faculdade de Medicina Veterinária de Lisboa, 1300-477 Lisboa, Portugal; Natural Resources Institute Finland (Luke), Latokartanonkaari 9, FI-00790 Helsinki, Finland; NordGen-Nordic Genetic Resources Centre, Ås, Norway; Natural Resources Institute Finland (Luke), Halolantie 31A, FI-71750 Maaninka, Finland; Arctic Centre, University of Lapland, Rovaniemi, Finland; Department of Agricultural Sciences, University of Helsinki, P.O. Box 28, FI-00014 Helsinki, Finland; CIBIO, Centro de Investigação em Biodiversidade e Recursos Genéticos, InBIO Laboratório Associado, Campus de Vairão, Universidade do Porto, 4485-661 Vairão, Portugal

## Abstract

**Background:** The drastic change in global climate has led to in-depth studies of the genetic resources of native cattle adapted to challenging environments. Native cattle breed data may harbor unique variants that will enable the generation of new tools to improve the adaptation potential of commercial cattle breeds. Adipose tissues are key factors in the regulation of metabolism and energy balance and are crucial for the molecular switches needed to adapt to rapid environmental and nutritional changes. The transcriptome landscape of four adipose tissues was used in this study to investigate the effect of the environment on the gene expression profiles of three local breeds, Yakutian cattle (Sakha Republic), Northern Finncattle (Finland), Mirandesa cattle (Portugal) and commercial Holstein cattle.

**Results:** A total of 26 animals (12 cows, 14 bulls) yielded 81 samples of perirenal adipose tissue (n=26), metacarpal adipose tissue (n=26), tailhead adipose tissue (n=26) and prescapular adipose tissue (n=3). More than 17,000 genes were expressed in our dataset. Principal component analysis of the normalized expression profiles revealed a differential expression profile of the metacarpal adipose tissue. We found that the genes upregulated in the metacarpal adipose tissue of Yakutian cattle, such as *NR4A3*, *TEKT3*, and *FGGY*, were associated with energy metabolism and response to cold temperatures. In Mirandesa cattle, the upregulated genes in perirenal adipose tissue were related to immune response and inflammation (*AVPR2, CCN1*, and *IL6*), while in Northern Finncattle, the upregulated genes appeared to be involved in various physiological processes, including energy metabolism (*IGFBP2*). According to the sex-based comparisons, the most interesting result was the upregulation of the *TPRG1 gene* in three tissues of Yakutian cattle females, suggesting that adaptation is related to feed efficiency.

**Conclusions:** The highest number of differentially expressed genes was found between Yakutian cattle and Holstein, several of which were associated with immunity in Yakutian cattle, indicating potential differences in disease resistance and immunity between the two breeds. This study highlights the vast difference in gene expression profiles in adipose tissues between breeds from different climatic environments, most likely highlighting selective pressure and the potential significance of the uniquely important regulatory functions of metacarpal adipose tissue.

## Background

In domestic cattle studies, whole-genome sequencing has been successfully applied to investigate genetic diversity, genomic architecture, history of cattle populations, selection signatures, and economically and physiologically important genomic variations. These are all important investigations for characterizing cattle genetic resources for agriculture and food production [1,2]. However, thus far, genomic characterization at the functional level is lacking for native cattle breeds, which is crucial to understanding their ability to adapt to various environmental circumstances [3]. RNA sequencing may allow us to unravel the critical knowledge needed for understanding how environmental circumstances, demographic factors and breeding histories are reflected in gene expression profiles in different cattle breeds [3]. Moreover, transcriptome profiling can reveal important candidate genes for production, fertility and health traits in domestic animal species and breeds and the genetic and evolutionary basis of these complex traits [4–6].

Here, we used high-throughput RNA sequencing technology to investigate and compare gene expression profiles in four adipose tissues (metacarpal, perirenal, tailhead and prescapular adipose tissues), representing visceral, peripheral and bone marrow fat tissue, in three native breeds and one commercial cattle breed. These adipose tissues are important organs for many physiological functions for survival and successful reproduction [7]. Due to the energy storing and channelling properties of the adipocytes, adipose tissue plays a crucial role in energy metabolism, for example, cold-induced adaptive thermogenesis, as well as in cushioning internal organs and insulating the body [8]. Adipose tissues help control energy balance and metabolic activity and are also capable of restructuring on the basis of nutritional changes [8]. There are two main types of adipose tissues in mammals: brown and white.

White adipose tissue stores energy, whereas brown adipose tissue dissipates stored energy as heat by burning fatty acids to maintain body temperature [9]. Fat in bone marrow acts as an energy reservoir contributing to metabolic processes and undergoes dynamic changes, for example, as a result of starvation [10]. Weldenegodguad et al.[9] recently reported that the gene expression profiles of metacarpal tissues were distinct from those of perirenal and prescapular fats in semi-domestic reindeer (*Rangifer tarandus*), providing interesting insights into the nature of adipose tissues, with candidate genes involved in immune response and energy metabolism.

The native cattle breeds included in this study have adapted to various biogeographical and production environments and have different genetic origins and breeding histories— Mirandesa cattle from Portugal, Northern Finncattle from Finland and Yakutian cattle from the Sakha Republic (Yakutia), the Russian Federation. In addition, we investigated the Holstein cattle breed, which is the high-producing and most popular dairy cattle breed worldwide (Figure 1) [3,9]. Studying the genetic resources of native, locally adapted cattle is essential due to climate change; the current productive commercial breeds, such as the Holstein, may not express the genetic variations important for future breeding and adaptation to changing environments [11]. Yakutian cattle, for example, live in the area of Sakha (Yakutia), which has some of the most extreme low temperatures in the settled world (-70°C to +30°C); thus, this breed is of great interest when studying adaptation to cold [2,9]. Adaptation to cold temperatures simultaneously implies adaptation to a different type of diet, which in this case consists of wild forage.

**Fig.1.**
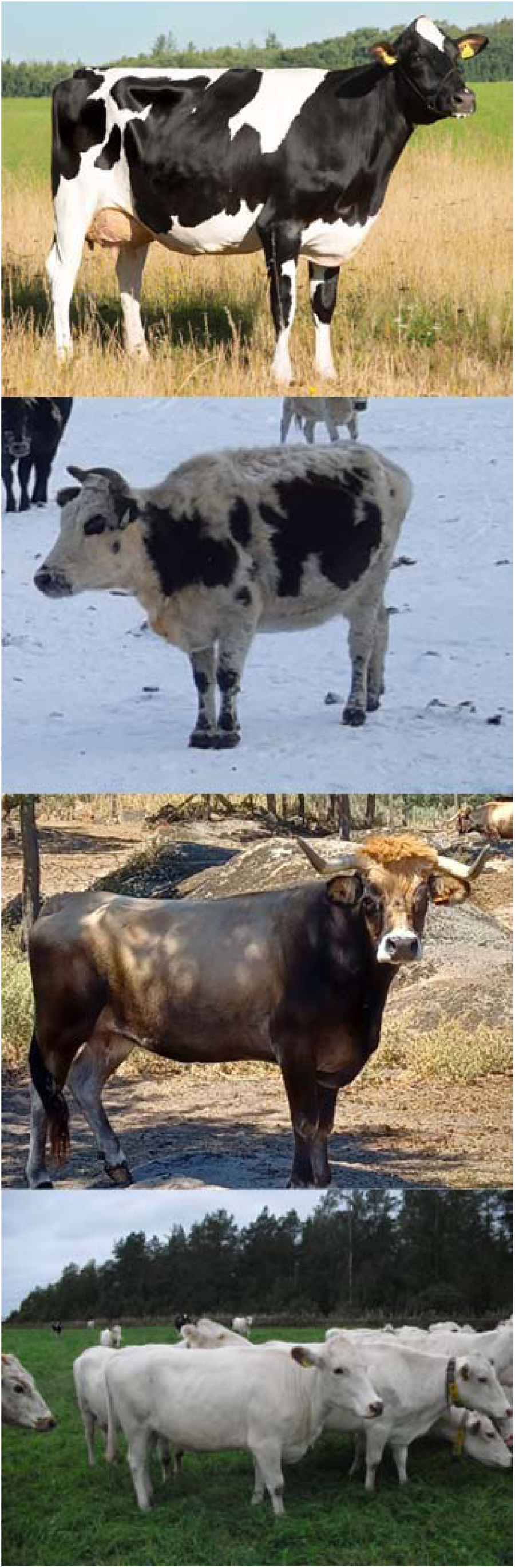
The breeds analysed in this study (top to bottom: Holstein cattle, Yakutian cattle, Mirandesa cattle and Northern Finncattle). Photos: Finnish Animal Breeding Association (Holstein cattle), Juha Kantanen (Yakutian cattle, Mirandesa cattle, Northern Finncattle).

Northern Finncattle exhibit certain similarities: the breed is well adapted to harsh northern environments and grazing, is also cultivated on natural pastures and is used for milk and meat production [2,13]. Challenging environmental conditions and limited natural resources have had an impact on the traditional diet of Northern Finncattle. Their feed typically included cooked soft stewed mash made from ingredients such as lichen and other less nutritious forage [14]. On the other hand, the Mirandesa cattle breed resides in southern Portugal and is characterized by a moderate-to-warm maritime climate. Previous studies have highlighted the unique nature of Mirandesa cattle, as it is one of the most genetically distinct Portuguese breeds and has low genetic diversity [15]. This breed is known for its excellent meat quality, even marbling fat, longevity, fertility, temperament and adaptation to limited rough forage and the environment.

Given the unique nature of the aforementioned breeds, it is important to study gene expression in adipose tissues to understand the physiological importance of these genes for the adaptative capacity of native breeds of cattle in the northern and southern Eurasian environments. Moreover, we are interested in identifying differences in gene expression profiles between intensively selected international Holstein cattle and locally adapted native breeds.

## Methods

### Sample collection

A total of 81 adipose tissue samples were collected from 12 adult cows and 14 bulls (2–7 years old) of native breeds from different geographical locations and climates, namely, Northern Finncattle from Finland, Mirandesa cattle from Portugal, Yakutian cattle from Sakha in the Russian Federation and Holstein cattle from Finland (Fig. 1; Table 1). The samples were randomly collected at slaughter from northern and central Finland, northern Portugal and northern and central Sakha (the Eveno-Bytantay and Magan regions, Sakha, Yakutia, the Russian Federation) during October 2015 and March 2016. Three different types of adipose tissue were collected: perirenal adipose tissue around the kidneys, tailhead adipose tissue between the tailhead and the tuber ischii, and metacarpal adipose tissue from the bone marrow in the diaphysis of the metacarpal bone (left front leg) (Supplementary Fig. 1). In addition, three prescapular adipose tissue samples were collected from Yakutian cattle beneath the cervical muscle in front of the scapula. The samples were stored in RNAlater® Solution (Ambion/QIAGEN, Valencia, CA, USA) after collection. Among the Yakutian samples, 3 were derived from castrated males and 3 from uncastrated males, whereas all the males from Finland (n = 5, Northern Finncattle) and Portugal (n = 3, Mirandesa) were uncastrated. All protocols and sample collections were performed in accordance with the legislations approved by the Russian authorization board (FS/UVN 03/163733/07.04.2016) and the Animal Experiment Board in Finland (ESAVI/7034/04.10.07.2015).

**Table 1.** Sample summary.

| **Breed** | **MAT (F+M)** | **PAT (F+M)** | **TAT (F+M)** | **PSAT (F+M)** |
| --- | --- | --- | --- | --- |
| **Holstein** | 3+0 | 3+0 | 3+0 | 0+0 |
| **Northern Finncattle** | 3+5 | 3+5 | 3+5 | 0+0 |
| **Mirandesa** | 3+3 | 3+3 | 3+2 | 0+0 |
| **Yakutian** | 3+6* | 3+6* | 3+6* | 0+3 |

Metacarpal adipose tissue = MAT, perirenal adipose tissue = PAT, tailhead adipose tissue = TAT and prescapular adipose tissue = PSAT. A total of 81 samples representing female (F) and male (M) individuals were sequenced. For instance, 3+5 refers to 3 females and 5 males. Out of the 6 Yakutian males, denoted by an asterisk “\*”, 3 were castrated and 3 were noncastrated.

### RNA extraction and sequencing

RNA extraction, library preparation, and sequencing were performed at The Finnish Functional Genomic Center (FFGC), Turku, Finland. Total RNA was extracted from adipose tissues (ca 30 mg/sample) using a Qiagen AllPrep DNA/RNA/miRNA kit. RNA extractions were performed according to the manufacturer’s protocol. The quality of the extracted RNA was confirmed with an Agilent Bioanalyzer 2100 (Agilent Technologies, Waldbronn, Germany), and the concentration of each sample was measured with a Nanodrop ND-2000 (Thermo Scientific, Wilmington, USA) and a Qbit(R) Fluorometric Quantification Kit (Life Technologies). All the samples had an RNA integrity number (RIN) above 7.5.

Library preparation was performed according to the Illumina TruSeq® Stranded mRNA Sample Preparation Guide (part #15031047). Unique Illumina TruSeq indexing adapters were ligated to each sample to pool several samples later in one flow cell lane. Library quality was inferred with an Advanced Analytical Fragment Analyser, and library concentration was inferred with a Qubit fluorometer; only good-quality libraries were sequenced.

The samples were normalized and pooled for automated cluster preparation, which was carried out with the Illumina cBot station. The libraries were analysed on an Illumina HiSeq 3000 platform. Paired-end sequencing with a 2 × 75 bp read length was performed with an 8 + 8 bp dual index run. Base calling and adapter trimming were performed using Illumina’s standard bcl2fastq2 software.

### Sequence data analysis

The overall quality of the raw RNA-seq reads in fastq and aligned reads in BAM format were examined using FastQC v0.11.7 [16]. The FastQC reports were summarized together with the results from other analyses using MultiQC v.1.7 [17]. We used Spliced Transcripts Alignment to a Reference (STAR) (version 2.7.8a) [18] for sequence alignment. Prior to alignment, a StarGenome was created using star v 2.7.8a with the parameters *--runThreadN 4, --runMode genomeGenerate, and --sjdbOverhang 74* using the cattle reference genome (ARS-UCD1.2) and transcriptome (ARS-UCD1.2.103.gtf), which were downloaded from Ensembl. The same version of STAR was also used for alignments with the following parameters: *--readFilesCommand zcat, --runThreadN 4, --outSAMtype BAM SortedByCoordinate, --quantMode GeneCounts, --twopassMode Basic.* Quality control checks of the resulting BAM files resulting from STAR alignment were performed with RSeQC v4.00 [19]. Gene body coverage showed uniform coverage, and junction saturation ranged from 107,569 to 323,347 at 100% of the reads. RSEM v 1.3.3 [20] quantification was performed to establish gene expression levels for RNA-seq data with the parameter *–paired-end*.

We used DESeq2 1.32.0 [21] to perform gene expression analysis for tissues, gender and breeds. We implemented differential gene expression analysis between breeds by incorporating gender as a second factor because we have clearly noted that gender does have an effect on gene expression. There were no male Holstein cattle; thus, separate gene expression analysis was performed between commercial Holstein females and females of the native breeds. Lowly expressed transcripts (<= 5 reads) were filtered out prior to running the DEseq command. We performed principal component analysis (PCA) using PlotPCA to obtain an overview of the sample distribution. Differentially expressed genes (DEGs) were identified by using several pairwise comparisons within and between populations. We used Log2Foldchange (log2FoldChange > 1.49) and adjusted P-value (padj < 0.05) to select the list of DEGs. The additional gene information, including the gene name and gene description, was retrieved for all DEGs using the biomaRt 2.48.3 bioconductor package [22]. Volcano plots were generated using the EnchancedVolcano R package. Finally, we performed Gene Ontology (GO) and Kyoto Encyclopedia of Genes and Genomes (KEGG) pathway analyses on all tissue-specific uniquely expressed genes and later DEGs using Gage v3.16 [23] for GO and KEGG analyses using biological pathway datasets with the parameter *same.dir=TRUE*.

## Results

### RNA-Seq and mapping

A total of 2,830,100,766 read pairs were generated from data of 81 samples of perirenal (n=26), metacarpal (n=26), tailhead (n=26) and prescapular (n=3) adipose tissues. After sequencing, a total of 1156 multi-lane Fastq files were merged into 162 forward and reverse files representing 81 samples. The Phred quality scores from the reads of all the samples were > 30 (30.57-40.11). The number of reads per sample ranged between 27.1 million and 63.2 million, with an average of 35.2 million (see Additional File 1:S1).

### Gene expression overview

A total of 20,714 genes (of 27,607 total listed bovine genes) were expressed in 81 samples. According to the RNA-Seq Expectation Maximization (RSEM) results, the highest number of genes with an abundance > 0.1 transcripts per million reads (TPM) were expressed in metacarpal adipose tissue (n=17,401), followed by tailhead adipose tissue (n=16,992), perirenal adipose tissue (n=16,802) and prescapular adipose tissue (n=16,060). Principal component analysis (PCA) of the normalized expression profiles revealed that PC1 and PC2 together explained ∼50% of the variance. PC1 alone explained 44% of the variance and clearly separated metacarpal adipose tissue from the other types of adipose tissue (Fig. 2B). There were far fewer obvious divisions between the rest of the tissues, with tailhead and perirenal adipose tissues overlapping remarkably.

**Fig.2.**
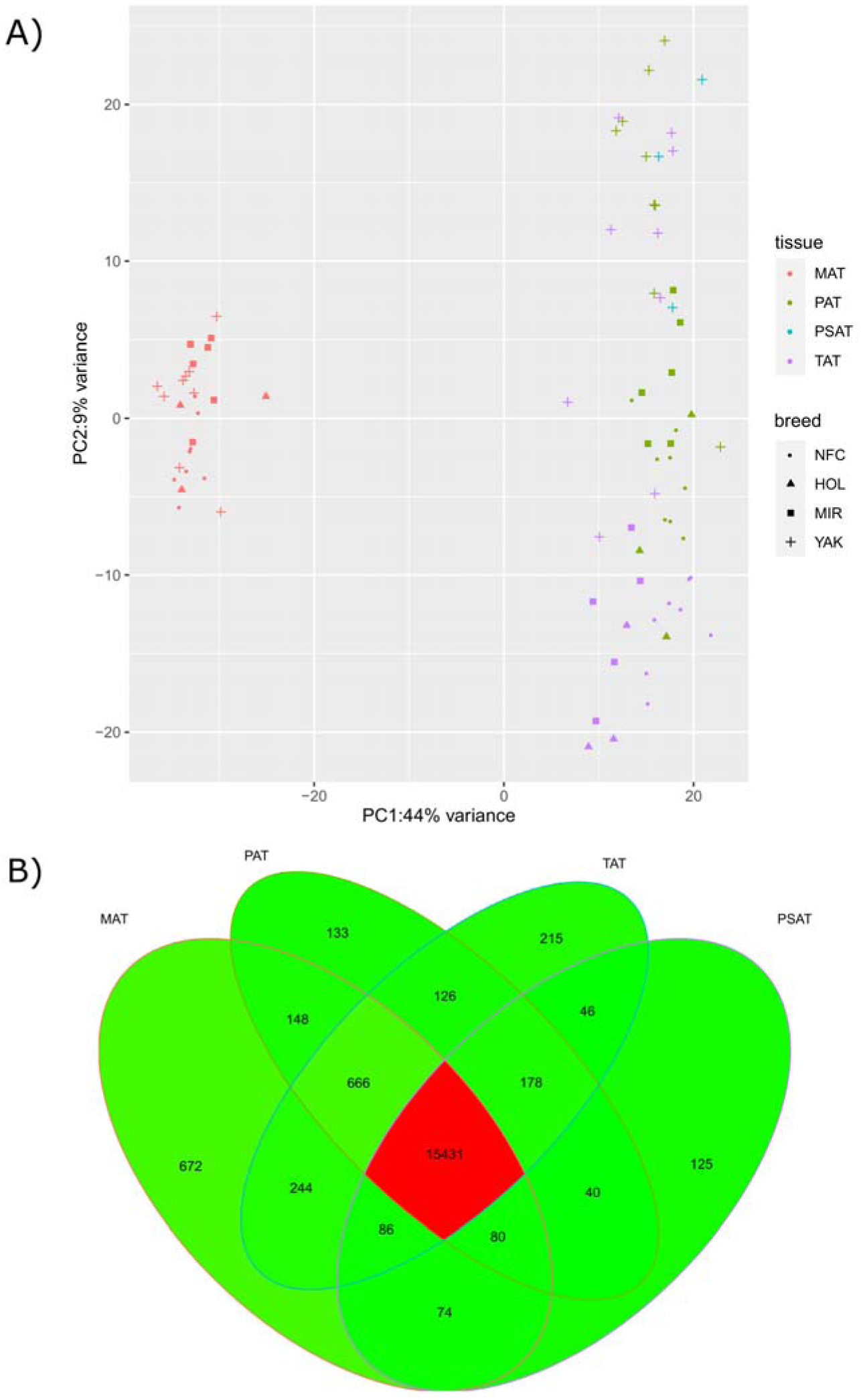
Graphical summary of gene expression in adipose tissues. A) PCA plot based on the gene expression of the tissue samples representing Yakutian cattle (YAK), Mirandesa cattle (MIR), Holstein cattle (HOL) and Northern Finncattle (NFC) samples based on expression profiles, with dot colours indicating tissue and breed; B) shared and unique genes expressed in the tissues: a Venn diagram of expressed genes with abundance >0.1 transcripts per million (TPM) in metacarpal adipose tissue (MAT), tailhead adipose tissue (TAT), perirenal adipose tissue (PAT) and prescapular adipose tissue (PSAT).

As shown in Fig 2A, 15,431 genes were shared by all four tissues. The metacarpal adipose tissue (n=672; Additional File 3:S1) harbored the highest number of uniquely expressed genes, followed by the tailhead (n=215; Additional File 3:S2), perirenal adipose tissue (n=133; Additional File 3:S3) and prescapular adipose tissue (n=125; Additional File 3:S4).

In metacarpal adipose tissue, the HOX family of genes (*HOXD13*, *HOXD12*), olfactory receptor genes (e.g., *OR10AG82P, OR2AD1, OR2AZ1, OR2J1*), and solute carrier family genes (e.g., *SLC12A3, SLC25A2, SLC26A9*) were uniquely expressed. The top GO terms associated with uniquely expressed genes in metacarpal adipose tissue (Additional file 4:S1) included “GO:0048513 animal organ development”, “GO:0048522 positive regulation of cellular process”, “GO:0051716 cellular response to stimulus”, “GO:0048518 positive regulation of biological process”, “GO:0050896 response to stimulus”, “GO:0016043 cellular component organization”, “GO:0071840 cellular component organization or biogenesis”, “GO:0048856 anatomical structure development as well as metabolic process, organic substance metabolic process, cellular metabolic process, primary metabolic process, and macromolecule metabolic process”. Interestingly, one of the GO terms in this list was reproduction, which may indicate the importance of metacarpal adipose tissue in cellular development, metabolism and reproduction. Moreover, two KEGG pathways (metabolic pathways and neuroactive ligandlJreceptor interaction) were associated with uniquely expressed genes in metacarpal adipose tissue (Additional file 4:S2).

The most abundant gene in the tailhead adipose tissue (Additional file 3:S2) was *bta-mir-192,* which is a microRNA possibly associated with genes targeting lipogenesis and the regulation of adipose deposition and differentiation [24]. Genes such as *HOXC12*, *HOXC13*, *SERPINA3-1* and *SERPINA3-3* were uniquely expressed in tailhead adipose tissue, and *SERPINA3-1* may be associated with growth traits and development in Chinese cattle [25]. Similarly, uniquely expressed genes in perirenal adipose tissue (Additional File 3:S3), a set of microRNAs, including *mir-197* and *TEX11,* are associated with andrological growth traits, specifically testicular size and circumference [26,27] (Additional File 3:S3). Finally, the uniquely expressed genes in prescapular adipose tissue included *PHGR1*, *KRT7* and *DRGX* (Additional File 3:S4). We did not find any GO terms or KEGG pathways for the lists of the highest abundance genes in tailhead, perirenal or prescapular adipose tissues.

### Castrated vs. uncastrated male samples

As indicated in Table 1, we obtained both castrated and noncastrated male samples from Yakutian cattle. Previous adipose tissue studies have shown gene expression differences in the adipose tissues of castrated and noncastrated mammals; for example, in castrated mice, brown adipose tissue will begin to convert to white adipose tissue [28]; therefore, it was important to compare the gene expression profiles of adipose tissue samples between the castrated and noncastrated male samples of Yakutian cattle. However, only a few DEGs were detected between Yakutian cattle castrated and noncastrated male samples. Only one DEG (*TDH*) was detected in the metacarpal adipose tissue (Additional File 2:S1), and this gene was upregulated in the noncastrated samples. Similarly, four DEGs were present in the tailhead (Additional File 2:S1) adipose tissue, of which two (*C11H2orf50 and ENSBTAG00000007075*) were upregulated in the castrated animals, and two (*SARDH and GSTA2*) were upregulated in the noncastrated animals. The expression of all three DEGs (*ENSBTAG00000025258, CXCL9, and ENSBTAG00000052522*) in perirenal adipose tissue was upregulated in the castrated animals (Additional File 2:S1). Owing to the minimal effect of castration in the present Yakutian cattle samples, all the samples were grouped together in further analyses, taking these DEGs into consideration in our conclusions (See Additional File 2:S1).

### Gender differences

Our comparison of the gene expression profiles between male and female individuals of the native cattle breeds revealed a total of 345 significant DEGs in the three adipose tissues (Table 4; Additional files 10, 11, and 12). The highest number of DEGs between the sexes was identified in the tailhead adipose tissue.

In metacarpal adipose tissue from Yakutian cattle, there were 26 significant DEGs between the male and female samples. Among Yakutian cattle females, the DEG with the highest log2foldchange (2.55) was *TPRG1,* which is associated with high weight gain traits in beef cattle with low feed intake [29]. Among Yakutian cattle males, the DEG with the highest log2fold change was *GDF5, which is* associated with body traits in both *Bos taurus* and *Bos indicus* [30]. In Mirandesa cattle females, upregulated genes were associated with latent tuberculosis (*CD209*) [31], ovarian morphology and milk traits (*ADCY5*) [43], mastitis and mastitis immunity (*OSMR*, *PTX3*) [44,45] and temperament (*BARHL2*) [32]. In male Mirandesa cattle, the top upregulated genes were linked to immune function (*SLC7A8*) [33] as well as susceptibility to BSE (PRND) and BVD (DHCR24) [34,35]. Similarly, the most upregulated genes in Northern Finncattle females and males were *GNAO1 and CACNA1G,* respectively*. GNAO1 is* known to affect lactation traits [36], and *CACNA1G* is associated with feed efficiency [37].

Interestingly, in perirenal adipose tissue, the gene with the greatest upregulation in Yakutian cattle females was still *TPRG1,* highlighting the importance of this gene and the general trend toward feed efficiency in Yakutian cattle. The most highly upregulated gene in Yakutian cattle males was *IL20RA,* which is associated with susceptibility to bovine tuberculosis in cattle [38]. In Mirandesa cattle females, the highest known upregulated gene was *MT1E,* which is associated with postpartum oxidative stress [39]. Moreover, this gene was also present in the breed comparisons. This was followed by *ACKR1*, *MT2A*, and *S100A12,* which are associated with mastitis and mastitis response [40–42]. The highest known upregulated gene in Northern Finncattle females was *JCHAIN,* associated with mastitis response [43]. This was followed by *UCHL1* and *DYNC1I1,* which are associated with splayed forelimbs at birth and limb development as well as body conformation traits [44–46].

In tailhead adipose tissue, the most upregulated DEG in Yakutian cattle females was *GLP1R,* which is associated with energy efficiency and a reduction in feed intake [47], which correlates with the trend in Yakutian cattle females. The most upregulated gene in Mirandesa cattle females was *MT1E,* which could also indicate the calving ability of the breed.

### Differential gene expression profiles in native cattle breeds

The analysis of DEGs between three native breeds was conducted by incorporating sex as a second factor. The highest number of DEGs (n=26) was found in tailhead adipose tissue between Mirandesa cattle and Northern Finncattle (Table 2, Additional File 7:S3).

**Table 2.** Number of upregulated genes in males and females.

| Breed/Tissue | Northern Finncattle |  | Mirandesa cattle |  | Yakutian cattle |  |
| --- | --- | --- | --- | --- | --- | --- |
|  | Male | Female | Male | Female | Male | Female |
| **MAT** | 11 | 36 | 7 | 13 | 20 | 6 |
| **PAT** | 12 | 35 | 7 | 7 | 8 | 4 |
| **TAT** | 39 | 59 | 27 | 41 | 9 | 4 |

MAT: metacarpal adipose tissue, PAT: perirenal adipose tissue, TAT: tailhead adipose tissue

In metacarpal adipose tissue, the comparison of Yakutian and Mirandesa cattle yielded 17 DEGs, with 7 upregulated for Yakutian and 10 upregulated for Mirandesa. The gene with the greatest upregulation in Yakutian cattle compared to Mirandesa cattle was *NR4A3*, which is associated with fat deposition and carbohydrate metabolism [48,49]. The most highly upregulated gene in Mirandesa was *PPP1R14C*, which is associated with immune tolerance and longevity in cattle [50,51]. A comparison of Yakutian cattle and Northern Finncattle yielded 7 genes, with 1 (*EML6*) gene upregulated in Yakutan cattle and 6 genes upregulated in Northern Finncattle. In addition, *EML6* is associated with reproductive traits and body composition traits in cattle [52,53]. On the other hand, *CLDN4*, one of the most upregulated genes in Northern Finncattle, was found to be expressed in the rumen in association with lactation in live yeast supplementation of cattle [54] and in maintaining the integrity of epithelial tissues. Moreover, *CLDN4* has been found to be associated with various diseases, including cancer and inflammatory bowel disease. Finally, the comparison of Mirandesa cattle and Northern Finncattle cattle yielded only one gene. The upregulation of the *SLC38A2* gene in Mirandesa was associated with carcass traits and pregnancy maintenance [55,56].

In tailhead adipose tissue, the comparison of Yakutian and Mirandesa cattle yielded four genes in total; one (*JPH4*) was upregulated in Yakutian, and three (*PTHLH*, *ENSBTAG00000048635*, and *PPP1R14C*) were upregulated in Mirandesa. *JPH4* is known to be associated with feed efficiency in pigs [57]. For *PPP1R14C,* we observed a pattern of expression similar to that in metacarpal adipose tissue. A comparison of Yakutian cattle and Northern Finncattle yielded two genes, with *IGFBP2* upregulated in Yakutian cattle and associated with puberty, female reproduction and lactation and *SLC2A1* upregulated in Northern Finncattle associated with milk quality/yield and bovine respiratory disease. A comparison of tailhead adipose tissue from Mirandesa and Northern Finncattle revealed 26 genes, 19 of which were upregulated in Mirandesa cattle and seven of which were upregulated in Northern Finncattle. The most upregulated gene in Mirandesa cattle in this comparison was *IL6*, which is associated with susceptibility to paratuberculosis, mastitis and trypanosomiasis. In comparison with those in Mirandesa, the expression of the top upregulated genes in Northern Finncattle is associated with milk production [58], immunity and disease [59].

In perirenal adipose tissue, a total of nine genes were significantly differentially expressed between Yakutian and Mirandesa cattle, eight of which were upregulated in Mirandesa cattle. *PLEKHG7*, the only gene upregulated in Yakutian cattle, is associated with coat colour in Vrindavani cattle [60]. A comparison of Yakutian and Northern Finncattle revealed 17 DEGs, with 2 genes upregulated in Yakutian cattle and 15 genes upregulated in Northern Finncattle. The most upregulated gene in Yakutian cattle was *PRSS42*, which is associated with lipid metabolism during the follicular and luteal stages of the bovine estrous cycle [61] and plays a role in murine spermatogenesis [62]. Similarly, *EGR1*, one of the top upregulated genes in Northern Finncattle, is associated with oocyte maturation, adipocyte differentiation and adipogenesis [63,64]. A comparison of the Mirandesa and Northern Finncattle genomes yielded 4 genes. Only one gene, *MT1E*, was upregulated in the Mirandesa cattle. The Northern Finncattle had 3 upregulated genes of which *KIAA0513* is within the top 5% of Consistently Introgressed Windows of Interest in Selection (CIWIS) in the Chianina, Romagnola, and Marchigiana breeds [65] and is associated with residual feed intake in indicine cattle [66].

### Differences in gene expression between the commercial and native breeds

The Holstein breed is the most popular dairy cattle worldwide and is selected for milk production traits [67]. Here, we examined whether there are differences in gene expression between commercial breeds and less selected local native breeds. In general, we identified many more DEGs in the comparisons between the native breeds and Holstein cattle (Table 3) than in the comparisons performed between only the native breeds (Table 2).

**Table 3.** Number of differentially expressed genes (DEGs) between commercial and native breeds according to female sample.

| Comparison | Total DEGs | Upregulated | Downregulated |
| --- | --- | --- | --- |
| **Metacarpal adipose tissue** |  |  |  |
| YAK vs HOL | 78 | 51 | 27 |
| NFC vs HOL | 10 | 8 | 2 |
| MIR vs HOL | 45 | 32 | 13 |
| **Tailhead adipose tissue** |  |  |  |
| YAK vs HOL | 580 | 357 | 223 |
| NFC vs HOL | 53 | 32 | 21 |
| MIR vs HOL | 142 | 79 | 63 |
| **Perirenal adipose tissue** |  |  |  |
| YAK vs HOL | 519 | 258 | 261 |
| NFC vs HOL | 82 | 52 | 30 |
| MIR vs HOL | 433 | 251 | 182 |

YAK: Yakutian cattle, MIR: Mirandesa cattle, NFC: Northern Finncattle

**Table 5.**
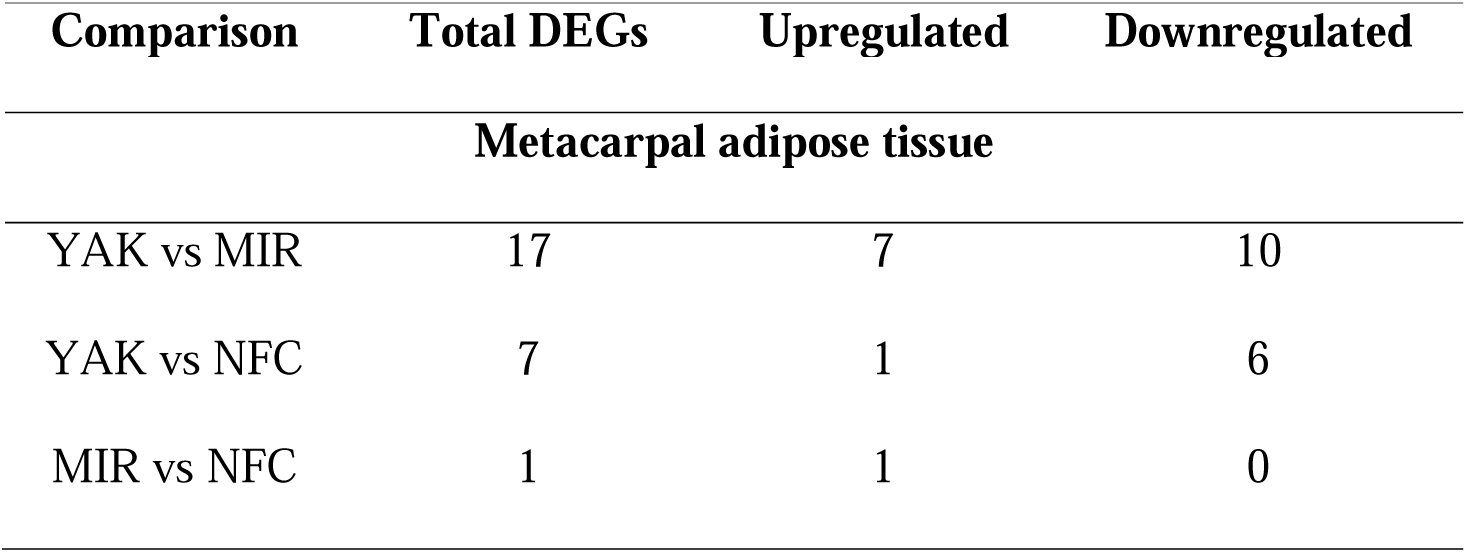
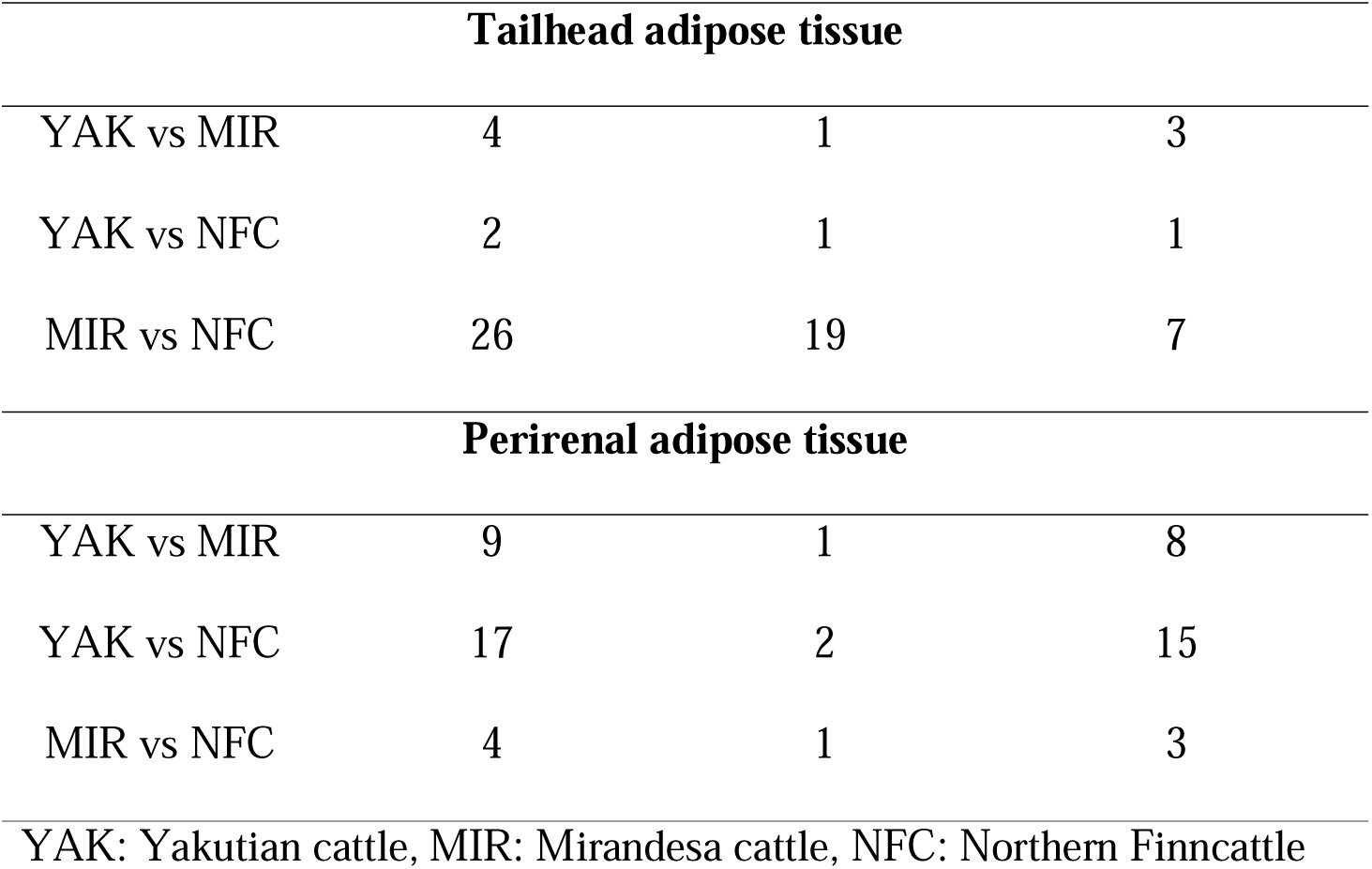
Number of differentially expressed genes (DEGs) between the native breeds.

| Comparison | Total DEGs | Upregulated | Downregulated |
| --- | --- | --- | --- |
| **Metacarpal adipose tissue** |  |  |  |
| YAK vs MIR | 17 | 7 | 10 |
| YAK vs NFC | 7 | 1 | 6 |
| MIR vs NFC | 1 | 1 | 0 |

| Tailhead adipose tissue |  |  |  |
| --- | --- | --- | --- |
| YAK vs MIR | 4 | 1 | 3 |
| YAK vs NFC | 2 | 1 | 1 |
| MIR vs NFC | 26 | 19 | 7 |
| Perirenal adipose tissue |  |  |  |
| YAK vs MIR | 9 | 1 | 8 |
| YAK vs NFC | 17 | 2 | 15 |
| MIR vs NFC | 4 | 1 | 3 |

YAK: Yakutian cattle, MIR: Mirandesa cattle, NFC: Northern Finncattle

Holstein cattle had the most differentially expressed genes when compared with the aboriginal Yakutian cattle in all three adipose tissues (Table 3). In the tailhead adipose tissue, 580 DEGs were identified in the Yakutian cattle vs. Holstein comparison, while in the perirenal adipose tissue, 519 DEGs were found, and in the metacarpal tissue, 78 DEGs were found. The expression of genes involved in the immune response and resistance/susceptibility to disease (*PRF1*, *CD8B*, *IDO1*, *CCL5*, *CD3D*, *CD3E*, and *GNLY*), in milk composition traits (*FCGR2B*) and in feed intake (*GIMAP4*) was strongly upregulated in Yakutian cattle (log2- fold change LFC >4). In Mirandesa, strongly upregulated genes were found in the following categories: fertility (*STC1*, *TCIM*), milk production/composition (*MAPK15*), meat quality (*CLK1*), feed efficiency (*MT1E*, *CCL8*, *HI-4*), and disease response (*HP*, *IL6*). Interestingly, the lowest number of DEGs in all analyses was found between the native Northern Finncattle and Holstein cattle. The genes highly upregulated in Northern Finncattle were associated with the following categories of meat quality: *PLTP*, *ACTN2,* and *ABCD2* [68,69]; immune response (*CDHR1*) [70]; and milk composition traits as well as thermotolerance in other species (*DUSP1*) [71,72].

GO enrichment analyses of the significant DEGs revealed a total of 126 GO terms with a p-value < 0.05 only for the three native breeds vs Holstein comparisons: Yakutian cattle in the tailhead adipose tissue, Yakutian cattle in the perirenal adipose tissue and Mirandesa cattle in the perirenal adipose tissue (Additional File 8:S10). The DEGs between Yakutian cattle and Holstein cattle in the tailhead adipose tissue had the highest number (n=86) of associated GO terms. There were 59 GO terms associated with upregulated genes in Yakutian cattle (Additional File 8:S10) among the top (p-value <0.05), such as “GO:0031328 positive regulation of biosynthetic process” and “GO:0009891 positive regulation of cellular biosynthetic process”. Other interesting GO terms in this list included “GO:0006955 immune response”, “GO:0006952 defense response”, “GO:0098542 defense response to other organism”, and “GO:0002682 regulation of immune system process”, which indicate differences in disease resistance and general immunity between Holstein cattle and Yakutian cattle. There were 27 GO terms associated with upregulated genes in Holstein cattle compared with Yakutian cattle (Additional File 8:S9), the top of which included “GO:1901564 organonitrogen compound metabolic process”, “GO:0008610 lipid biosynthetic process”, “GO:0071702 organic substance transport”, “GO:0046395 carboxylic acid catabolic process” and “GO:0071704 organic substance metabolic process”. Other interesting GO terms associated with upregulated genes in this category included “GO:0006629 lipid metabolic process”, “GO:0006631 fatty acid metabolic process”, and “GO:0044255 cellular lipid metabolic process”, which suggest a possible difference in lipogenesis between these two breeds.

The KEGG pathways associated with the genes upregulated in the Yakutian cattle vs Holstein cattle in the tailhead adipose tissue comparison included “Th1 and Th2 cell differentiation”, “Th17 cell differentiation”, “antigen processing and presentation”, “chemokine signaling pathway”, and “cytokinelJcytokine receptor interaction” (Additional File 9:S1). On the other hand, the KEGG pathways associated with downregulated genes in Holstein cattle in the tailhead adipose tissue were “metabolic pathways”, “carbon metabolism”, “valine, leucine and isoleucine degradation”, “oxidative phosphorylation” and “thermogenesis” (Additional File 9:S2).

A total of 30 GO terms were generated for the DEGs between Yakutian cattle and Holstein cattle in the perirenal adipose tissue. There were 5 GO terms associated with upregulated genes in Yakutian cattle, “GO:0002376 immune system process” and “GO:0006955 immune response” (Additional File 8:S12), again indicating possible differences in disease resistance between these two breeds. The downregulated genes in Holstein cattle were associated with 25 GO terms (Additional File 8: S11), such as “GO:0044281 small molecule metabolic process”, “GO:0044238 primary metabolic process”, “GO:0044237 cellular metabolic process” and “GO:0008152 metabolic process”, as well as “GO:0006629 lipid metabolic process” and “GO:0008610 lipid biosynthetic process”.

Pathway analysis revealed 18 KEGG pathways associated with upregulated genes in Yakutian cattle, for example, “viral protein interaction”, “metabolic pathways” and “thermogenesis”, while no KEGG pathways were found for downregulated genes in the perirenal tissue of Holstein cattle (Additional File 9:S3).

The significant DEGs between Mirandesa cattle and Holstein cattle in the perirenal adipose tissue generated 10 GO terms in total. There was one upregulated GO term, “immune system process” (Additional File 8:S14), and 9 downregulated GO terms, including “lipid metabolic processes” (Additional File 8:S13).

## Discussion

Transcriptome studies have shown that gene expression patterns in adipose tissue differ between breeds, sexes, and adipose tissue depots and, more recently, that there are differences in adaptation to extreme environments [9,73]. Here, we studied the transcriptome profiles of four adipose tissues from three native cattle breeds and one commercial breed, all of which originated from different geographical locations and climates. In the present study, we identified a total of 16,060–17,401 genes in the analysed tissues, which covered approximately 63.8% of the list of genes available for the *Bos taurus* reference genome (ARS-UCD1.2.). We found that metacarpal adipose tissue displayed a distinct pattern of gene expression compared to the other three tissues, which is in line with the functional role of this tissue, as highlighted in a recent reindeer study [9]. Similarly, metacarpal adipose tissue had the highest number of uniquely expressed genes (671), and functional annotation of those unique genes indicated the role of this tissue in the development and regulation of cellular processes as well as metabolism. Several genes belonging to the homeobox (HOX) family were uniquely expressed in metacarpal adipose tissue. In particular, *HOXD13* had the highest abundance among all uniquely expressed genes in the metacarpal adipose tissue (Additional File 3: S1). *HOXD13* is a highly conserved gene belonging to the HOX family of genes and is responsible for morphogenesis, limb development and genital development [74,75]. In murines, the *HOX* gene family has been associated with cell differentiation toward adipogenesis [76], and in cattle, the *HOX* gene family is associated with muscularity traits [77] and limb development [45]. The distinct gene expression profiles of metacarpal adipose tissue and the presence of homeobox genes among the uniquely expressed genes in metacarpal adipose tissue are in agreement with the findings of a recent study in reindeer (*Rangifer tarandus*) [9]. On the other hand, in tailhead tissue, the uniquely expressed members of the HOX family of genes are *HOXC12* and *HOXC13*, which have been previously associated with thermotolerance in African cattle [78].

Here we investigated the gene expression profiles of physiologically vital adipose tissues in native cattle breeds adapted to North European boreal (Northern Finncattle), South European Mediterranean (Mirandesa) and North Asian extreme continental (Yakutian cattle) environments. Moreover, the gene expression profiles of these native breeds were compared with those of the most common commercial cattle breed, Holstein, to detect possible human-generated selection effects from functional genomic and origin points of view. In our Yakutian cattle sample set, we also had RNA-Seq data from castrated males, but castration appeared to have a minimal effect on the gene expression profiles; thus, the data from the castrated and uncastrated males were pooled in the subsequent analyses. However, within the native breeds, we identified several DEGs between the female and male samples; therefore, sex was considered a second factor in the DEG analysis between the three native cattle breeds. The breedwise comparisons showed that Yakutian cattle exhibited superior disease resistance traits to those of Mirandesa and Northern Finncattle cattle, thus promoting adaptation to a challenging environment. Several of the genes downregulated in Yakutian cattle were related to the susceptibility of a host to diseases, including *Mycobacterium avium* paratuberculosis, bovine tuberculosis, mastitis, brucellosis, and bovine respiratory disease (*CD209*, *EEF1A2*, *SLC2A1*). The *CD209* gene plays a key role in the pathogenesis of bovine paratuberculosis; furthermore, there is evidence for a genetic basis for susceptibility to *Mycobacterium avium* subspecies of bovine paratuberculosis in cattle [79]. This gene was downregulated in the metacarpal adipose tissue of Yakutian cattle in comparison to that of Mirandesa cattle. *ACKR1* has been shown to be related to mastitis, as it was found to be differentially expressed in comparison with healthy udder tissue in other studies [40]. This is interesting because local Yakutian cattle farmers ‘help’ their cows to stay resistant by exposing the udder to cold but also when it gets too cold, protecting the udders via thermal insulation. This also involves traditional culture-based observations as to which cows need such additional shelter and which are resilient enough to withstand this. We could hypothesize that this practice may have contributed to artificial selection to increase mastitis resistance. *SLC2A1* is associated with the immunological response to bovine respiratory disease in Washington and Colorado beef cattle. On the other hand, Yakutian cattle presented metabolism-related genes (e.g., *TPRG1* and *GLP1R*). Within-breed sex comparisons revealed that Yakutian cattle females exhibit superior metabolic characteristics to males. According to sex comparisons of all tissues of Yakutian cattle, *TPRG1* was highly upregulated. This gene is associated with feed efficiency traits, specifically high weight gain under low-feed conditions in cattle [29]. In tailhead adipose tissue, *GLP1R* was upregulated in females. In addition to being associated with feed efficiency, this gene is also associated with obesity in humans [80], and *GLP-1* analogues such as semaglutide are used for the treatment of obesity and diabetes [81]. These cattle do not eat industrially produced food. They feed of course in the summer on fresh grass, but in spring, they also eat some green plants from previous years on the shores of rivers. Moreover, in the late summer/autumn, they also travel to the forest and feed on shrubs, leaves and twigs. According to herders, this makes the animals more resistant to different kinds of feed. In the most important ‘bottleneck’ season for feeding, in spring, the feed that herders give to their animals is most different by sex: lactating females obtain the freshest hay. Herders try to ensure that they always receive green hay, while males obtain different qualities of hay. For example, hay from later harvests or of lower quality that was already brown. This practice, according to herders, is a regular measure in spring when good quality fodder is scarce, so they prioritize females that feed calves and people with milk. During fieldwork when one sample animal for this study was slaughtered in the village of Kustur, the local veterinarian measured the length of the intestines of the slaughtered animal and commented that it was much longer than that of other breeds of cattle in other areas. According to him, this is a result of the long-term adaptation of the animals to a more diverse diet and the need for the animals to digest “tougher” fodder such as twigs, branches and shrubs in the forest.

Similarly, the expression of genes related to lactation and other mammary traits (*SLC38A2*, *SLC35B1*), fertility (*SLC38A2*, *MT1E*) and feed intake/efficiency (*MT1E*, *SLC38A2*) was upregulated in Mirandesa cattle. This tracks the characteristics of this breed, which is well adapted to a rough forage environment. In addition, this breed is known for its ease of calving and quality of life. *SLC35B1* is associated with glucose transportation and the lactose biosynthesis pathway. This gene was found to be expressed in lactose synthesis [82].

*SLC38A2* has been found to be associated with postmortem carcass traits in Nellore cattle, specifically with the ribeye area and the amount of meat in the carcass [55]. This gene has also been associated with pregnancy maintenance in cattle [56] and peak lactation in sows [83]. This gene specifically plays an important role in amino acid transport during lactation in sows and may influence dairy quality. Interestingly, *MT1E* was found to be upregulated in Mirandesa cattle in all comparisons with other native breeds in perirenal tissue. This protein- coding gene is thought to enable zinc ion binding activity; zinc ions have a limited ability to bind to metallothionein, which is sensitive to oxidative stress. This oxidative stress results in elevated concentrations of free zinc and induces a pro-oxidative state. Therefore, this gene is associated with postpartum oxidative stress [39] in cattle and was also found to be associated with high residual feed intake in heifers adjusted for backfat thickness [84]. The same gene was found to be highly upregulated in all tissues of Mirandesa cattle females compared with males, possibly indicating that this feature is superior in females.

Based on the comparisons with Holstein cattle, it can be concluded that there are significant differences in gene expression between Holstein and Yakutian cattle in adipose tissues. The most differentially expressed genes were observed in tailhead adipose tissue from Holstein cattle compared to Yakutian cattle, followed by perirenal adipose tissue. Furthermore, GO term analysis of the DEGs revealed multiple upregulated immune-related terms in both tailhead and perirenal adipose tissues, indicating potential differences in disease resistance and immunity between the two breeds. Interestingly, the *MT1E* gene was again highly upregulated in Mirandesa cattle compared to Holstein cattle. The frequency of upregulation of this gene in Mirandesa cattle points to its importance in this breed’s traits. *DUSP1* was upregulated in Northern Finncattle and was previously associated with thermal tolerance in zebrafish; this gene regulates thermal limits through maintenance of mitochondrial integrity [71]. This finding points to the cold adaptive qualities of this native breed, which has been reared for centuries in northern Finland and as far north as the Lapland.

Additionally, KEGG pathway analysis of tailhead adipose tissues from Holstein cattle and Yakutian cattle suggested that there may be differences in energy metabolism and immune system functions between the two breeds. In perirenal adipose tissue, relatively fewer genes were differentially expressed between Holstein and Yakutian cattle, but these differences were still significant. GO term analysis revealed upregulation of immune system processes and immune responses in Holstein cattle compared to Yakutian cattle, which is consistent with the results observed in tailhead adipose tissue.

Overall, the results suggest that there are significant differences in gene expression and biological pathways between Holstein and Yakutian cattle, particularly in terms of immune-related functions and energy metabolism. Further studies on the functional roles of these genes and pathways could provide insights into the underlying mechanisms of these differences and their potential implications for production and disease resistance in cattle.

### Conclusions

The novelty of this study lies in its comprehensive investigation of differential gene expression in multiple cattle breed adipose gene transcriptomes. By examining different adipose tissue types (metacarpal, perirenal, and tailhead) across various cattle breeds (Yakutian, Mirandesa, Northern Finncattle, and Holstein), this study provides unique insights into the genetic adaptations that have evolved in response to diverse environmental conditions and selective pressures. There are likely differences in the functions and adaptations of these adipose tissues among the different breeds of cattle. For example, the genes upregulated in Yakutian cattle in metacarpal adipose tissue and perirenal adipose tissue, such as *TPRG1* and *IL20RA*, suggest adaptations related to feed efficiency and susceptibility to tuberculosis, respectively. In contrast, the upregulated genes in Perirenal adipose tissue of Mirandesa cattle, such as *CD209* and *MT1E*, suggest adaptations related to tuberculosis susceptibility and postpartum oxidative stress, respectively. Furthermore, the study highlights the differences in adipose gene expression between males and females within each breed, which may provide insights into the underlying mechanisms of these differences and their potential implications for production, disease resistance, reproductive traits and metabolism. Overall, this research provides valuable information about the physiological and metabolic differences between cattle breeds, which could lead to the identification of genetic markers associated with specific traits. However, further research is needed to fully understand the functions and mechanisms of the genes and pathways identified in the different adipose tissues. By identifying breed-specific differences in gene expression related to immune functions, energy metabolism, and adaptation to local environments, this research highlights the potential for using genetic markers to improve livestock management, breeding strategies, and disease resistance.

## List of abbreviations

DEG: Differentially expressed gene
YAK: Yakutian Cattle
MIR: Mirandesa Cattle
HOL: Holstein Cattle
NFC: Northern Finncattle
RIN: RNA integrity number
TPM: transcripts per million
MAT: Metacarpal adipose tissue
PAT: Perirenal adipose tissue
TAT: Tailhead Adipose Tissue
PSAT: Prescapular adipose tissue
GO: Gene Ontology
KEGG: Kyoto Encyclopedia of Genes

## Declarations

### Ethics approval

All protocols and sample collections were performed in accordance with the legislations approved by the Russian authorization board (FS/UVN 03/163733/07.04.2016) and the Animal Experiment Board in Finland (ESAVI/7034/04.10.07.2015).

### Consent for publication

Not applicable

### Availability of data and materials

Raw sequence data will be publicly available in ENA under accession PRJEB71475.

## Authors’ contributions

DR performed quality control of the sequence data, analysed the data and drafted the manuscript. AA, MW, and KP assisted DR in performing the bioinformatic analyses. MH, HL, JP, PS, FS, CG and JK collected the samples for the study. AA, MW, MH, FS, PU, CG, JK and KP provided useful advice on study design and interpretation of results. JK and KP conceived the original idea of the study and supervised the project. All authors read and approved the final manuscript.

## Acknowledgements

The authors thank Tiina Reilas, Innokentyi Ammosov, Ruslan Popov and animal owners for the collaboration in collection of research materials. We thank CSC-IT Center for Science, Finland for computational resources. This study was supported by the Finnish Functional Genomics Centre, University of Turku and Åbo Akademi and Biocenter Finland.

